# Flexible, fast and selective genetic manipulation of the vertebrate CNS with misPiggy

**DOI:** 10.1101/481580

**Authors:** Michal Slezak, Filip de Vin, Yohei Shinmyo, Mykhailo Y. Batiuk, Melvin Y. Rincon, Carmen Menacho Pando, Johann Urschitz, Stefan Moisyadi, Frank Schnütgen, Hiroshi Kawasaki, Matthew G. Holt

**Affiliations:** VIB Center for Brain and Disease Research, Herestraat 49, Leuven 3000, Belgium; KU Leuven Department of Neuroscience, Herestraat 49, Leuven 3000, Belgium; Department of Medical Neuroscience, Graduate School of Medicine, Kanazawa University, Ishikawa, 920-1192, Japan; Institute for Biogenesis Research, University of Hawaii, 1960 East-West Rd. E-124, Honolulu, Hawaii, 96822, USA; Department of Medicine 2, University Hospital Frankfurt, Goethe University, Theodor Stern Kai 7, Frankfurt am Main, D60590, Germany; LOEWE Center for Cell and Gene Therapy, University Hospital Frankfurt, Goethe University, Theodor Stern Kai 7, Frankfurt am Main, D60590, Germany; FCI, Frankfurt Cancer Institute, University Hospital Frankfurt, Goethe University, Theodor Stern Kai 7, Frankfurt am Main, D60590, Germany; Leuven Brain Institute, Herestraat 49, Leuven 3000, Belgium; Slezak and Pando: BioMedX GmbH, BioMedX Innovation Center, Im Neuenheimer Feld 515, D69120 Heidelberg, Germany

## Abstract

*In utero* electroporation is increasingly used for manipulation of gene expression in the mammalian central nervous system. Here we created a transposon-based single plasmid system, that allows long-term conditional loss- and gain-of-function experiments across major CNS cell types. The system is widely applicable across species, allowing fast, flexible and cheap cell manipulation in previously non-transgenic species.

---

Loss- and gain-of-function experiments have proved critical to our understanding of how genes function in the vertebrate central nervous system (CNS). Classically, such experiments have used genetically engineered mouse lines, which entails significant investments in terms of time and cost. Viral vectors (generally based on adeno-associated virus (AAV) or lentivirus) offer an alternative solution, with simplicity of manufacture allowing comparatively higher experimental throughput. To date, they have been the only solution when targeting genes and cell types for which transgenic lines do not exist. However, viral vectors have a considerable drawback in terms of the transgene size that they can carry (<10 kb)^1^. Given the recent explosion in sequencing data available for mice and other vertebrates^2^, which will require functional validation, a simple and cost-effective tool for both loss- and gain-of-function experiments would tremendously accelerate our understanding of gene function in the mammalian CNS (in a manner analogous to the impact made by the RNAi collections available to *Drosophila* researchers^3^).

*In utero* electroporation (IUE) systems are based on the injection of simple and cheap DNA plasmids into the ventricle of the embryonic brain, with uptake into CNS progenitors lining the ventricles facilitated by application of electrical pulses to the tissue using electroporators^4^. Since its introduction, IUE has emerged as a powerful tool, with a wide variety of imaginative modifications allowing investigation of cell autonomous and non-cell autonomous aspects of cell migration, differentiation and function in the brain (including in conditions of pathology) (reviewed extensively by LoTurco and colleagues^5^). However, what remains missing is a simple one plasmid system, guaranteeing temporally controllable loss- and gain-of-function experiments in easily identifiable cell populations, with simultaneous expression of probes to assess cell morphology and/or physiology^1^.

To address this issue, we developed misPiggy: <u>m</u>ulti-function <u>i</u>nducible <u>s</u>ystem based on <u>piggy</u>Bac). misPiggy is a unique single plasmid system specifically designed for IUE that allows cheap, fast and flexible screening of gene function, across different CNS cell types at various developmental stages, in multiple mammalian species.

Previous work has shown that stable, long-term expression of transgenes across CNS cell types is possible when IUE is combined with the use of genome integrating, transposon-based systems, in order to offset the plasmid dilution that occurs as embryonic progenitors divide and differentiate during post-natal development^5^. We chose to build a plasmid based on the piggyBac system, due to its proven transposition activity in mammalian cells^6^ and ability to efficiently deliver large DNA fragments (up to the size of bacterial artificial chromosomes) to cells^7^, which would facilitate further engineering of the system.

Hence, we based our work on the previously reported p*mhy*GENIE-3 system, which is unique in carrying both a self-inactivating piggyBac transposase and transgene cassette on a single plasmid^6^ (described in Figure 1a). Basic testing of p*mhy*GENIE-3 derived plasmids in IUE was performed using expression of a strong fluorescent reporter, TdTomato (Figure 1a). Reporter expression was driven by a modified astrocyte-specific GFAP promoter (mGFAP) (Supplementary Figure 1a), as astrocytes are known to proliferate extensively during the first few weeks of post-natal development, and so are liable to loss of episomal plasmids, providing easy scoring of transposon activity^8^. Using a three-electrode electroporation configuration^9, 10^ to specifically target the cortex of mouse brain, we confirmed that the transgene delivered from p*mhy*GENIE-3 was expressed specifically in multiple astrocytes across cortical layers into early adulthood (TdTomato signal; Figure 1b), even when the size of the plasmid was effectively doubled (from 10.6 kb to 21.3 kb) by increasing the size of the transgene cassette carried (from 2.6 kb to 13.3 kb) (Supplementary Figure 2). Such a transgene cassette should be sufficient for the majority of biological applications. Crucially, the combined effects of plasmid delivery and transgene expression did not result in detectable tissue damage, immune response, or compromised cell function in the adult brain (Figure 1b, c; Supplementary Figure 3).

**Figure 1.**
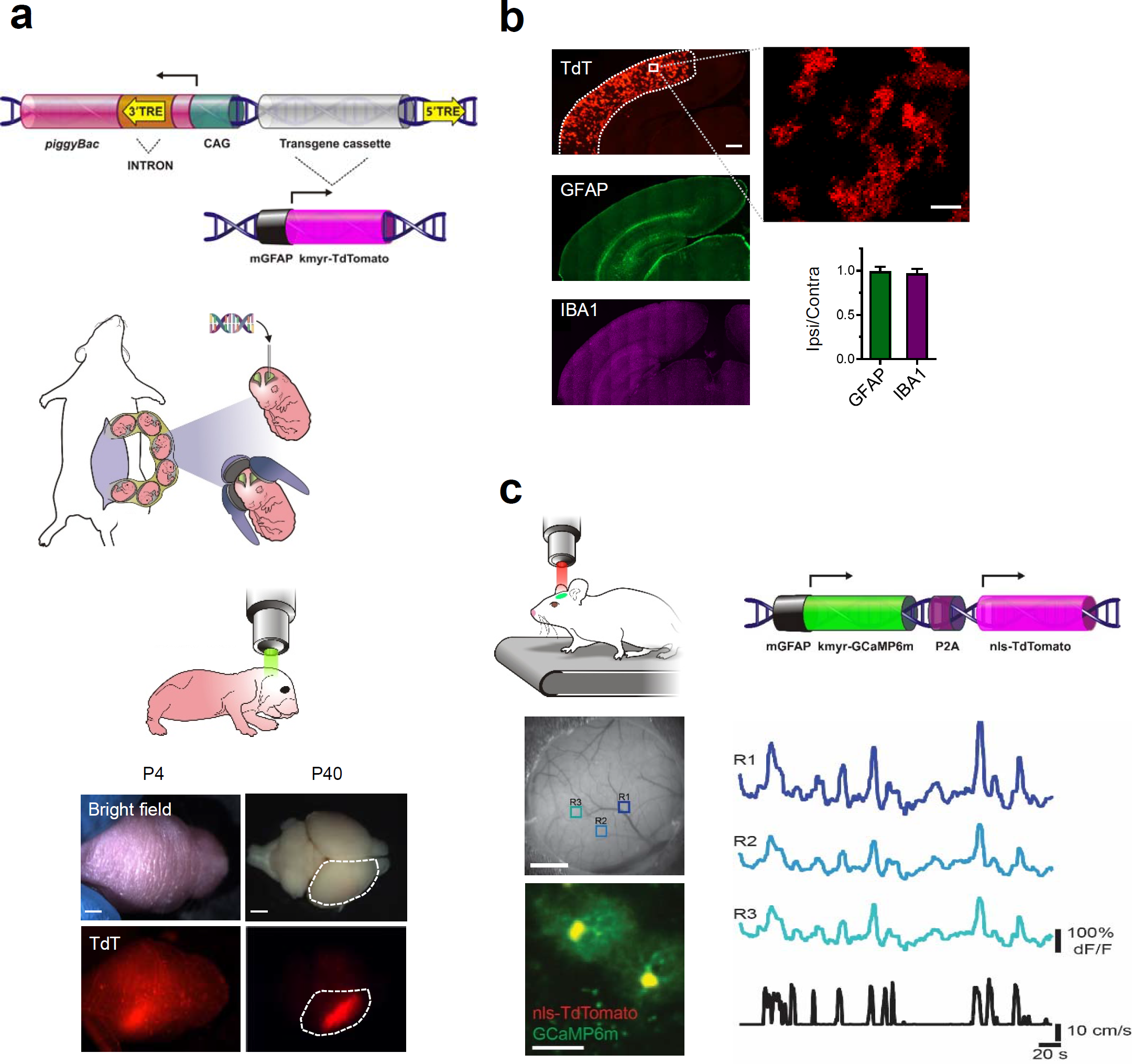
*In utero* electroporation using a single plasmid system gives efficient long-term gene expression in the mammalian CNS. (a) (b) Top: Schematic of p*mhy*GENIE-3. The transgene cassette is delineated by <u>T</u>erminal <u>R</u>epeat <u>E</u>lements (TREs) recognized by piggyBac transposase. Expression of piggyBac is driven by a CAG promoter, which is active in embryonic progenitors. Cell type specific promoters are incorporated into the transgene cassette. In this example, expression of membrane-tagged (kmyr) TdTomato (TdT) is driven by a modified GFAP promoter (mGFAP). Excision of the transgene cassette inactivates piggyBac expression limiting possible genotoxicity. Middle: p*mhy*GENIE-3 is microinjected into embryos and uptake facilitated by electrical pulses. A three-electrode system allows neural progenitors forming specific brain regions to be targeted. Bottom: TdT in cortex was detectable through the skull of mice at post-natal day 4 (P4) and was expressed into adulthood (P40). (b) TdT was expressed throughout the cortex on the electroporated side of the mouse brain. Cortex is delineated by white lines (low magnification). The mGFAP promoter limited expression to astrocytes (expanded). No CNS damage (increase in endogenous GFAP or increase in IBA1) was detected. Measurements were normalized to those made on the non-electroporated side. (c) Mice received a construct encoding the Ca^2+^ indicator GCaMP6m and nuclear-tagged TdT as a Ca^2+^-insensitive cell marker. Functional responses of astrocytes to locomotion^19^ were detected. Increases in GCaMP6m fluorescence in regions of interest (R1-R3) are plotted relative to baseline. Locomotion speed (black trace) (c). Scale bars, (a) 2 mm: (b) low magnification, 500 μm; high magnification, 30 μm: (c) cranial window, 1 mm; fluorescence imaging, 50 μm.

**Figure 2.**
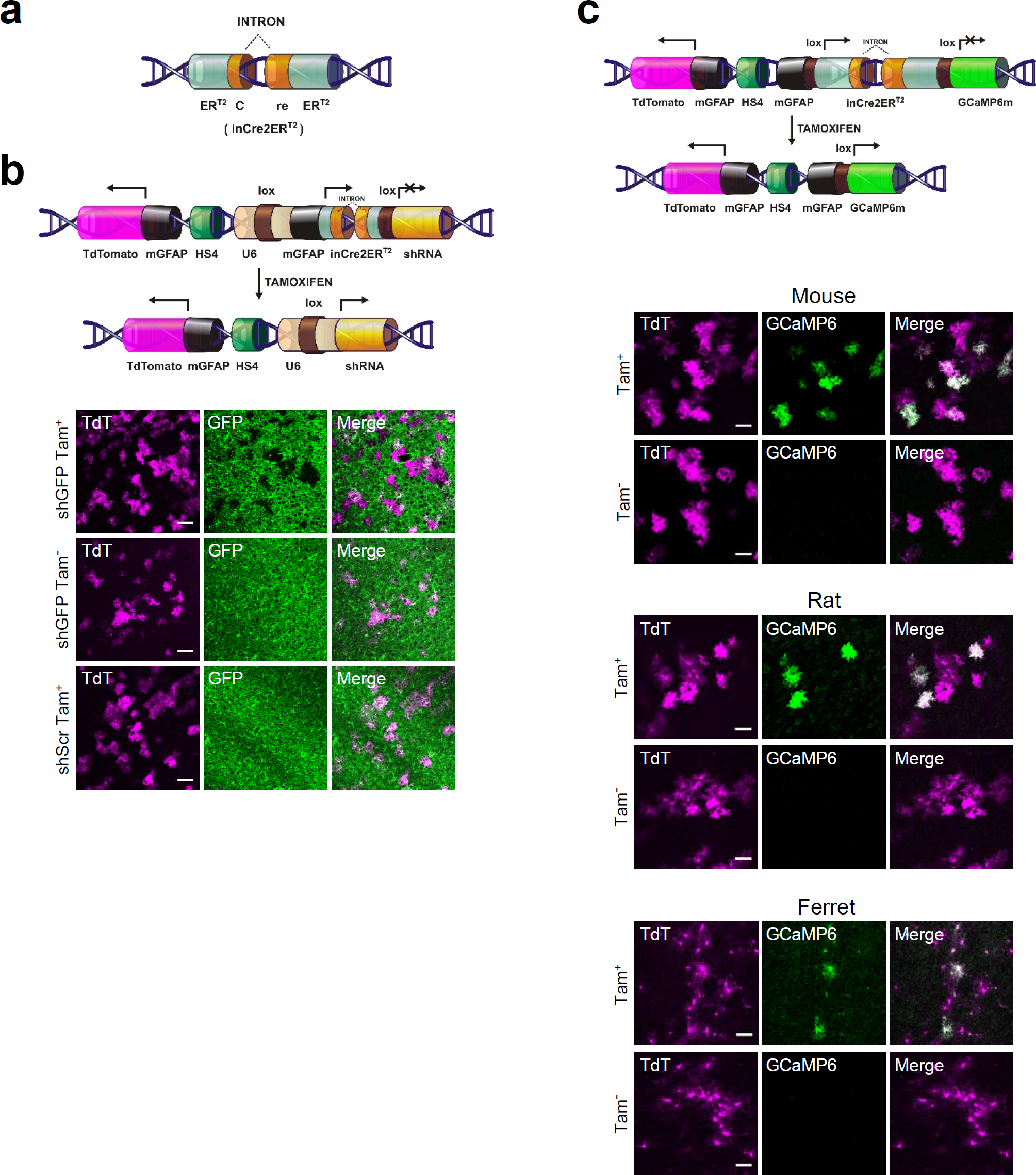
Cell type specific and inducible loss- and gain-of-function experiments from a single plasmid across multiple species. (a) Schematic representation of the tamoxifen inducible intronized Cre generated in this study. (b) Inducible loss-of-function. Representative images of brain slices obtained from adult Aldh1l1-EGFP mice (P21) electroporated at embryonic day 15.5 (E15.5). The schematic shows the key elements in the construct used. A mGFAP promoter was used to drive expression of TdT in astrocytes. Expression of tamoxifen inducible Cre in astrocytes was achieved using an independent mGFAP promoter. Following tamoxifen induction, recombination at loxP sites resulted in excision of Cre and reconstitution of an active U6 promoter driving shRNA expression. Animals were electroporated with a construct containing shRNA against GFP (shGFP) and treated with tamoxifen (Tam^+^) or vehicle (Tam^−^), or a construct containing a scrambled shRNA (shScr) (Tam^+^). Scale bars, 100 μm. (c) Inducible gain-of-function across mammalian species. The schematic shows the key elements in the construct used. Constitutive expression of TdT in astrocytes was driven by a mGFAP promoter. Tamoxifen inducible Cre was also expressed specifically in astrocytes but was under the control of an independent mGFAP promoter. Representative images of brain slices obtained from adult mice (P21) electroporated at E15.5. Following tamoxifen induction, recombination at loxP sites led to excision of Cre and the expression of the calcium indicator GCaMP6m. Rat and ferret were electroporated with the same construct at E16.5 and E33.5, respectively. Images show brain slices produced from animals treated either with vehicle alone (Tam^−^) or with tamoxifen (Tam^+^). Images show representative results. Scale bars, 50 μm.

These results confirm that a basic IUE system built on p*mhy*GENIE-3 largely matches or exceeds the performance of current systems. Therefore, we set out to engineer the backbone to incorporate extra elements in the form of easily modifiable ‘expression modules’ (see Supplementary Table 1). These modules allow simultaneous loss- and gain-of-function experiments in a cell type specific and temporally controlled manner and form the basis of misPiggy.

We chose to work with a variant of the well characterized and widely used Cre/lox system^11^, as a useful screening tool requires stable and irreversible genetic modification. Unfortunately, when Cre and loxP sites are contained on the same plasmid, this can lead to spontaneous recombination during cloning and propagation of the plasmid in bacteria^12^. To overcome this issue, we engineered a Cre containing a unique intron (inCre). This Cre variant is not expressed in prokaryotic cells during plasmid preparation but is actively expressed in eukaryotic cells. Strict ligand inducibility required the fusion of tamoxifen response elements (ER^T2^) to the N- and C-terminal ends of inCre (inCre2ER^T2^) (Figure 2a, Supplementary Figure 4). By placing the coding sequence for inCre2ER^T2^ under control of an appropriate promoter, between loxP sites, we generated a self-excising Cre variant, which allows temporal control of transgene expression in a cell type specific manner.

In order to perform cell type specific loss-of-function experiments, we decided to use short hairpin RNAs (shRNAs) for gene silencing, as these possess several advantages for rapid and flexible screening of gene function *in vivo*, based on issues such as cost, ease of use, time to phenotype and potential applicability across species (discussed extensively in^13^ and Supplemental Methods). Furthermore, extensive experience with the system means that there is an extensive collection of specific and validated shRNAs for the majority of (mouse) genes (https://www.ncbi.nlm.nih.gov/probe). Our system was designed around the split U6 promoter^14^, in which Cre-mediated recombination at TATA-lox sites leads to reconstitution of an active U6 promoter, which is capable of driving shRNA expression at high levels across all major cell types. To assess the potential for shRNA-mediated knock down, we performed IUE in transgenic mice expressing EGFP specifically in astrocytes (Aldh1l1-EGFP line). The construct used expressed both TdTomato (as a positive marker for electroporation) and inCre2ER^T2^ in astrocytes, under the control of independent mGFAP promoters. The construct also carried a validated shRNA against EGFP. Upon tamoxifen administration, cells in which recombination had occurred showed expression of TdTomato and silencing of EGFP expression, as judged by immunohistochemistry and flow cytometry (Figure 2b, Supplemental Figure 4).

In a further series of experiments, we tested the compatibility of the inCre2ER^T2^ system with protein overexpression. Initially, experiments were based on the conditional expression of the genetically encoded Ca^2+^ indicator, GCaMP6m, in astrocytes using the mGFAP promoter (Figure, 2c). Using this simple set up to score expression phenotype, we also extended the system for use in multiple cells types by testing the neuron specific human synapsin 1 (hSYN1) and oligodendrocyte specific proteolipid protein (PLP) promoters (Supplementary Figure 1b, 1c). Interestingly, in our hands, strict promoter specificity was dependent on the use of an inducible system, perhaps relating to protein perdurance after transient promoter activity in precursors.

Crucially, all manipulations were solely dependent on elements carried on the plasmid used for IUE and were entirely independent of the genetic background of the mouse used. This led us to explore the general applicability of our system across mammals. Hence, we explored whether misPiggy can be applied to other species amenable to IUE, such as rat^9^ and ferret^15^. These species are used in areas such as neurodevelopment, systems neuroscience, neuropharmacology and behavior, where differences in anatomy, biochemistry and physiology means they more accurately mirror specific aspects of human CNS development, function and disease, such as cortical folding. However, their use is generally limited in scope by a lack of tools for easy and robust genetic manipulation of the CNS. To test the cross-species applicability of our system, rats and ferrets were electroporated with the construct shown in Figure 2c. TdTomato was expressed constitutively in all species tested. Expression of GCaMP6 was strictly tamoxifen dependent (although the degree of recombination may have been impacted by the use of a non-optimized tamoxifen dose in both species). Interestingly, the mGFAP promoter used seemed highly specific for astrocytes in both rat and ferret (greater than 90% as based on cell morphology). However, absolute promoter specificity likely needs to be refined across species to accurately target specific cells types (which will depend on increasingly high resolution genomic information). These results demonstrate that the key components of our system retain functionality across multiple vertebrate species^16^.

In summary, we have developed a novel one plasmid-based toolbox, misPiggy, for long-term expression of multiple transgenes following IUE. Consistent with other IUE systems, the proportion of cells targeted is dependent on the size of the plasmid electroporated and can be adjusted by varying the amount of plasmid injected, the developmental stage at which electroporation is performed and the electroporation protocol (Supplementary Figure 2). Promoter specificity and strength, as well as recombination efficiency (which is related to the tamoxifen induction protocol used), also affect the performance of the system (data not shown). These parameters can be systematically adjusted according to experimental needs.

In contrast to other IUE based systems, however, the modular design of misPiggy, facilitated by the large cargo capacity of transposon-based systems, allows multiple expression modules to be used, allowing simultaneous loss- and gain-of-function experiments in defined cells. Furthermore, misPiggy is applicable across CNS cell types and, when combined with three-electrode electroporation, can target defined brain regions^10^. Such capacity dramatically increases the range of experiments possible with IUE and currently includes conditional rescue experiments following protein knock-down, but can also can be expanded to include the use of multi-promoter systems^1^ for effective targeting of the ever-expanding number of CNS cell subtypes identified using single cell approaches (for example using split-Cre approaches)^1^.

Modification of the individual expression modules to incorporate elements such as destabilized proteins^17^ as recombination markers, or easy conversion for guide RNA and Cas9^13^ expression, will undoubtedly refine the system, according to specific experimental needs. Furthermore, construct insertion (and copy number) can be specifically targeted to unique genomic sites by tethering piggyBac to a custom transcription activator like effector DNA-binding domain^18^. Finally, the re-engineered Cre allows experiments to be performed on an otherwise wild-type background, obviating the need for expensive transgenic lines, opening up the possibility of rapid and cost-effective genetic manipulation in species previously regarded as ‘non-transgenic’.

## Methods

Detailed methods are available in the online version of the paper.

## Supporting information

## Author contributions

MGH conceived and directed the project. JU and SM produced and provided the p*mhy*GENIE-3 plasmid used as the backbone in all experiments. FS developed the original inCre system modified in the paper. MS designed and cloned the majority of misPiggy plasmids, established the IUE protocol and performed *in vivo* imaging experiments. FdV cloned some constructs and performed the majority of immunohistochemistry experiments. He performed the IUE for rat experiments and, in addition, took care of all animal husbandry related to experiments with mice and rats. YS and HK performed all experiments on ferret. MYB performed flow cytometry experiments. MYR and CMP assisted with immunohistochemistry experiments. MGH wrote the final manuscript, with input from all other authors. All authors approved submission.

## Acknowledgements

MGH grateful acknowledges support from the European Research Council (Starting Grant 281961) and Fonds Wetenschappelijk Onderzoek (FWO) (Grants G066715N, G088415N and 1523014N). MS was the recipient of a Marie Curie Intra-European Fellowship (331018). SM and JU receive financial support from an NIH COBRE grant P20GM103457. MYR is a postdoctoral researcher with the FWO (133722/1204517N) and acknowledges the continuous support of the Fundación Cardiovascular de Colombia. FS is supported by individual grants from the Deutsche Forschungsgemeinschaft (SCHN1166/4-1) and from the Loewe Center for Cell and Gene Therapy. HK is supported by a Kanazawa University SAKIGAKE project 2018 and the Grants-in-Aid for Scientific Research program from MEXT, Japan.

The authors wish to express thanks to Sedef Dalbeyler, Arno Vandebroek, Jessica Bouhuijzen-Wenger, Steffen Kandler and Jeason Haughton for help and advice at various stages of this project. The authors also thank Dr. Vincent Bonin for access to his surgical facilities, 2-photon microscope and custom MATLAB scripts for image analysis. Drs. Tilmann Achsel and Carlos Dotti gave critical feedback on the manuscript. Schematic illustrations of the various plasmids were made by David Pennington (@ penningtonArt.co.uk).

## References

1. Luo, L., Callaway, E.M. & Svoboda, K. Genetic dissection of neural circuits. Neuron 57, 634–660 (2008).

2. Meadows, J.R.S. & Lindblad-Toh, K. Dissecting evolution and disease using comparative vertebrate genomics. Nat Rev Genet 18, 624–636 (2017).

3. Dietzl, G. et al. A genome-wide transgenic RNAi library for conditional gene inactivation in Drosophila. Nature 448, 151–156 (2007).

4. Bullmann, T., Arendt, T., Frey, U. & Hanashima, C. A transportable, inexpensive electroporator for in utero electroporation. Dev Growth Differ (2015).

5. LoTurco, J., Manent, J.B. & Sidiqi, F. New and improved tools for in utero electroporation studies of developing cerebral cortex. Cereb Cortex 19 Suppl 1, i120–125 (2009).

6. Marh, J. et al. Hyperactive self-inactivating piggyBac for transposase-enhanced pronuclear microinjection transgenesis. Proc Natl Acad Sci U S A 109, 19184–19189 (2012).

7. Rostovskaya, M. et al. Transposon-mediated BAC transgenesis in human ES cells. Nucleic Acids Res 40, e150 (2012).

8. Ge, W.P., Miyawaki, A., Gage, F.H., Jan, Y.N. & Jan, L.Y. Local generation of glia is a major astrocyte source in postnatal cortex. Nature 484, 376–380 (2012).

9. Szczurkowska, J. et al. Targeted in vivo genetic manipulation of the mouse or rat brain by in utero electroporation with a triple-electrode probe. Nat Protoc 11, 399–412 (2016).

10. dal Maschio, M. et al. High-performance and site-directed in utero electroporation by a triple-electrode probe. Nat Commun 3, 960 (2012).

11. Matsuda, T. & Cepko, C.L. Controlled expression of transgenes introduced by in vivo electroporation. Proc Natl Acad Sci U S A 104, 1027–1032 (2007).

12. Kaczmarczyk, S.J. & Green, J.E. A single vector containing modified cre recombinase and LOX recombination sequences for inducible tissue-specific amplification of gene expression. Nucleic Acids Res 29, E56–56 (2001).

13. Boettcher, M. & McManus, M.T. Choosing the Right Tool for the Job: RNAi, TALEN, or CRISPR. Mol Cell 58, 575–585 (2015).

14. Ventura, A. et al. Cre-lox-regulated conditional RNA interference from transgenes. Proc Natl Acad Sci U S A 101, 10380–10385 (2004).

15. Kawasaki, H., Iwai, L. & Tanno, K. Rapid and efficient genetic manipulation of gyrencephalic carnivores using in utero electroporation. Mol Brain 5, 24 (2012).

16. Lu, Y., Lin, C. & Wang, X. PiggyBac transgenic strategies in the developing chicken spinal cord. Nucleic Acids Res 37, e141 (2009).

17. Corish, P. & Tyler-Smith, C. Attenuation of green fluorescent protein half-life in mammalian cells. Protein Eng 12, 1035–1040 (1999).

18. Owens, J.B. et al. Transcription activator like effector (TALE)-directed piggyBac transposition in human cells. Nucleic Acids Res 41, 9197–9207 (2013).

19. Paukert, M. et al. Norepinephrine controls astroglial responsiveness to local circuit activity. Neuron 82, 1263–1270 (2014).

