## Supplementary material for "Flexible, fast and selective genetic manipulation of the vertebrate CNS with misPiggy"

| | MODULE 1 | | INSULATOR | MODULE 2 | | |
|---|---|---|---|---|---|---|
| Construct | Promoter | Transgene | | Promoter | Inducible cassette | Transgene |
| MP1 | mGFAP | Kmyr-TdT | - | - | - | - |
| MP2 | mGFAP | nls-TdT | - | - | - | - |
| MP3 | mGFAP | nls-TdT GFAP 11kb | - | - | - | - |
| MP4 | mGFAP | Kmyr-GCaMP6-P2A-nls-TdT | | | | |
| MP5 | mGFAP | TdT | HS4 | U6lox | mGFAP inCre2ER$^{T2}$ | shRNA CS WPRE |
| MP6 | mGFAP | TdT | HS4 | U6 | - | shRNA CS WPRE |
| MP7 | mGFAP | TdT | HS4 | mGFAP | lox-inCre2ER$^{T2}$-lox | GCaMP6 |
| MP8 | mGFAP | TdT | HS4 | hSYN | lox-inCre2ER$^{T2}$-lox | GCaMP6 |
| MP9 | mGFAP | TdT | HS4 | PLP | lox-inCre2ER$^{T2}$-lox | GCaMP6 |

Slezak, de Vin et al. Supplementary Table 1
