## Supplementary figures and images for "Flexible, fast and selective genetic manipulation of the vertebrate CNS with misPiggy"

### Supplementary file 2

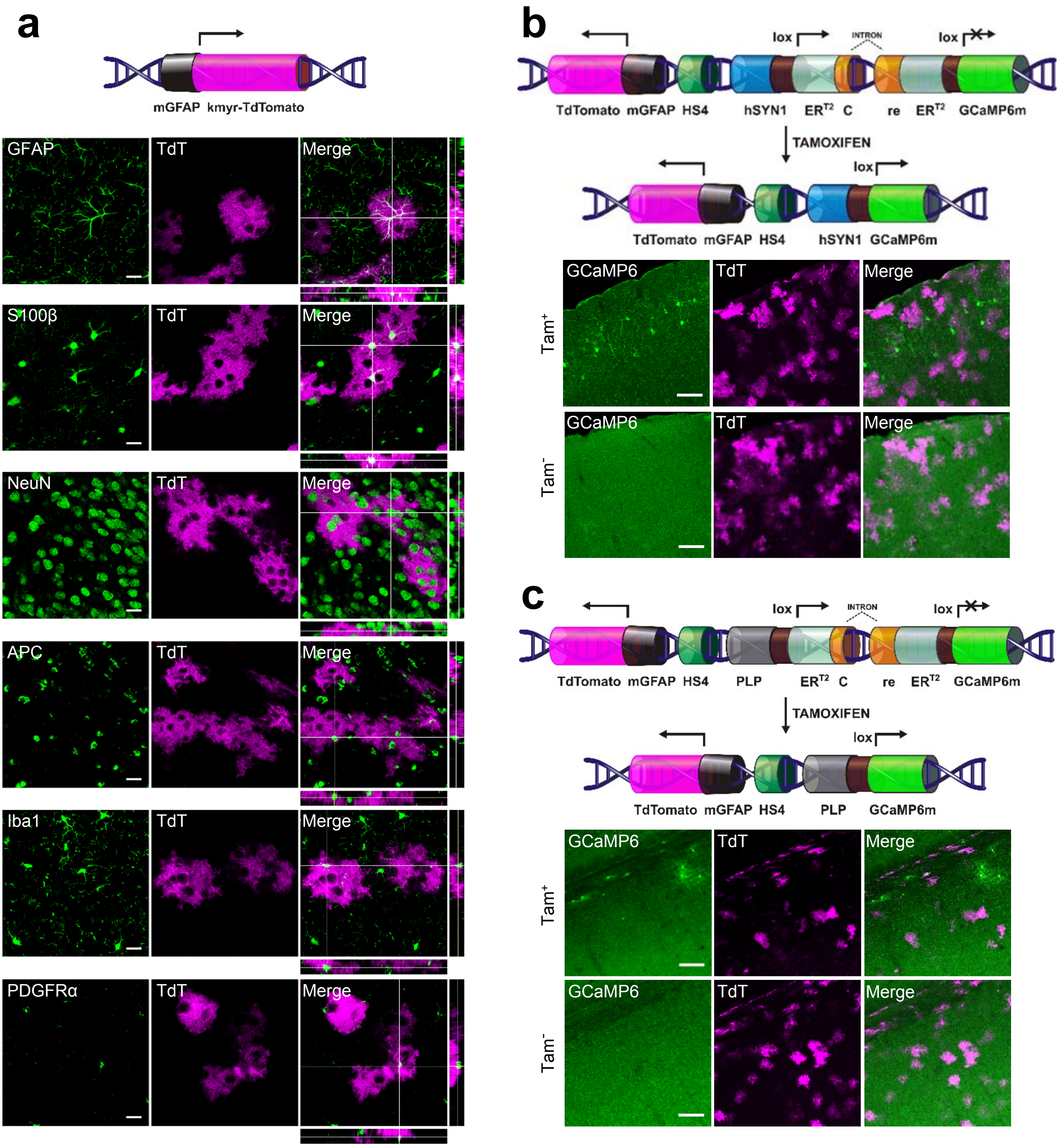


Slezak, de Vin et al., Supplementary Figure 1

### Supplementary file 3

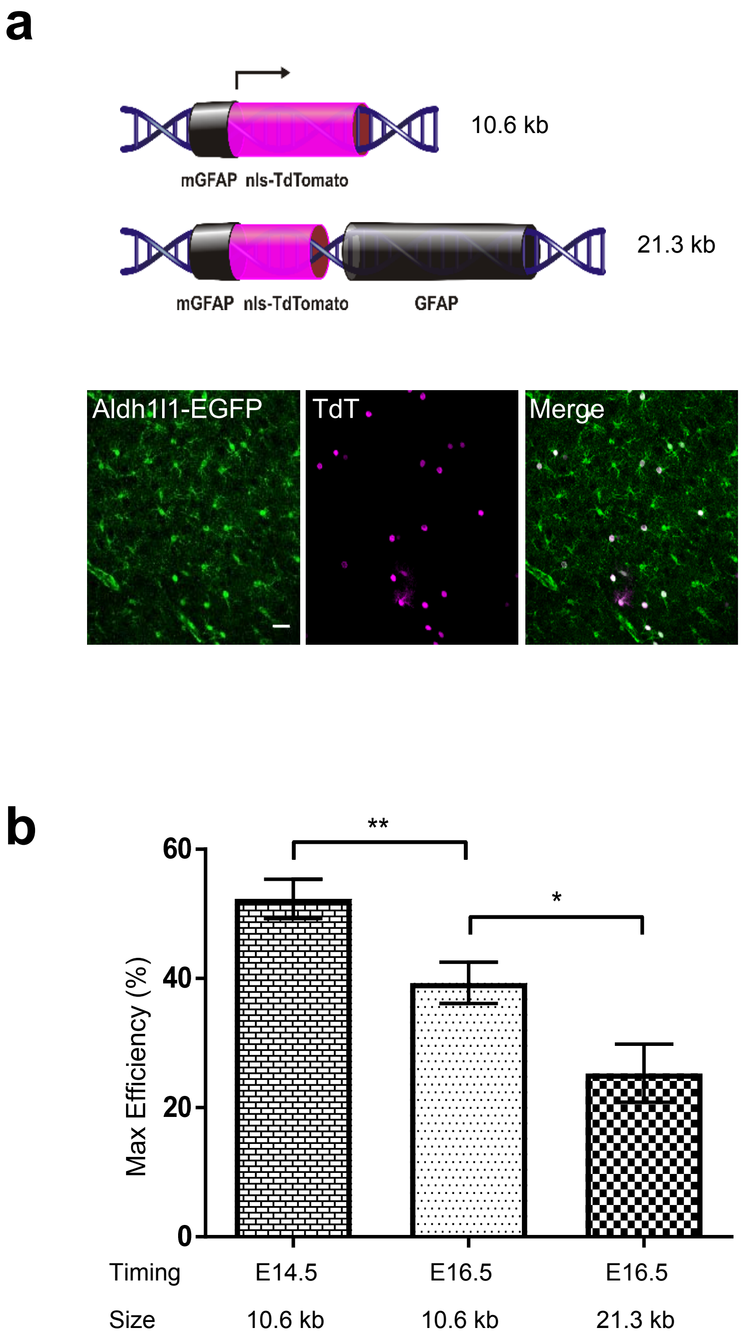


Slezak, de Vin et al., Supplementary Figure 2

### Supplementary file 4

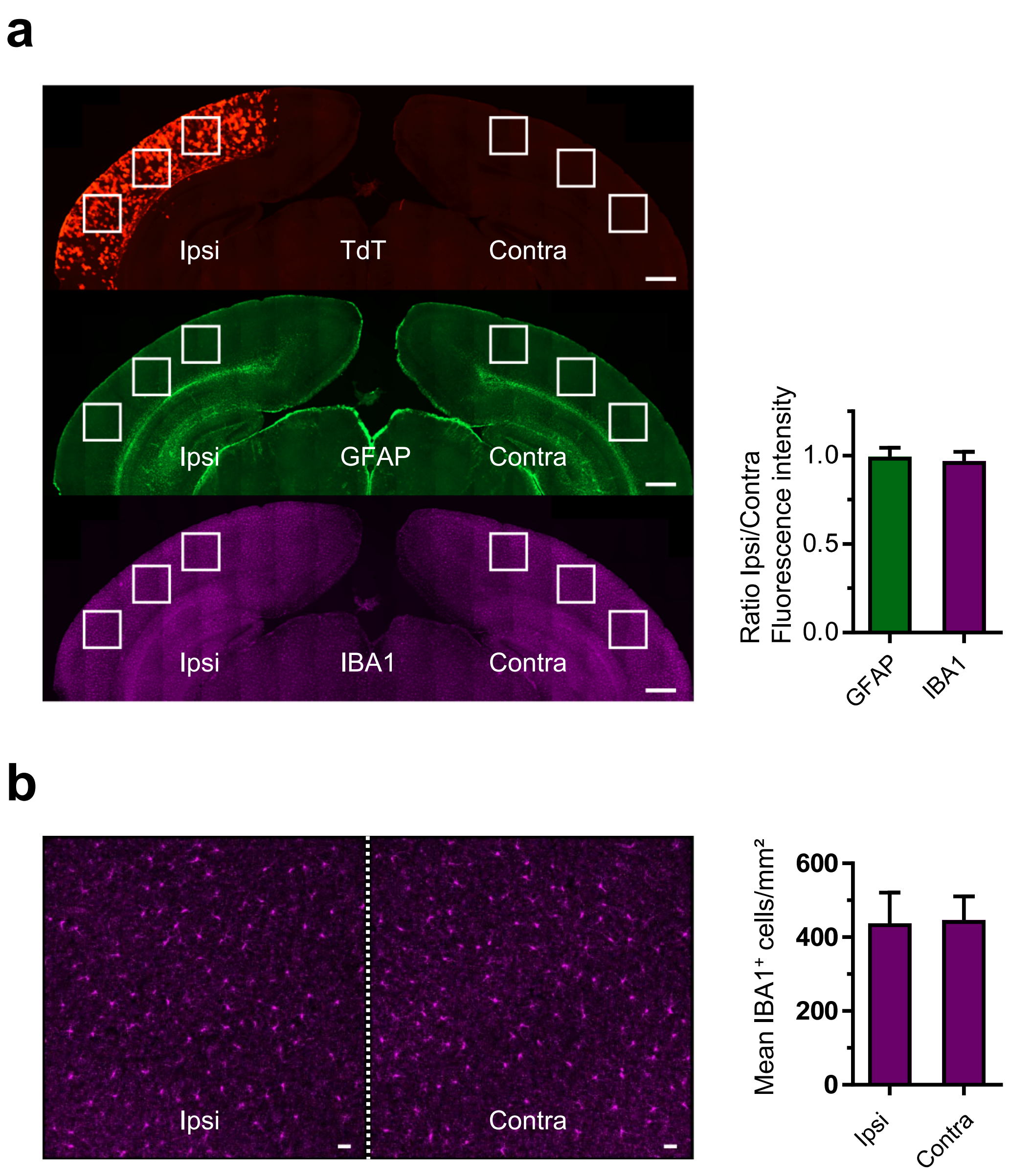


Slezak, de Vin et al., Supplementary Figure 3

### Supplementary file 5

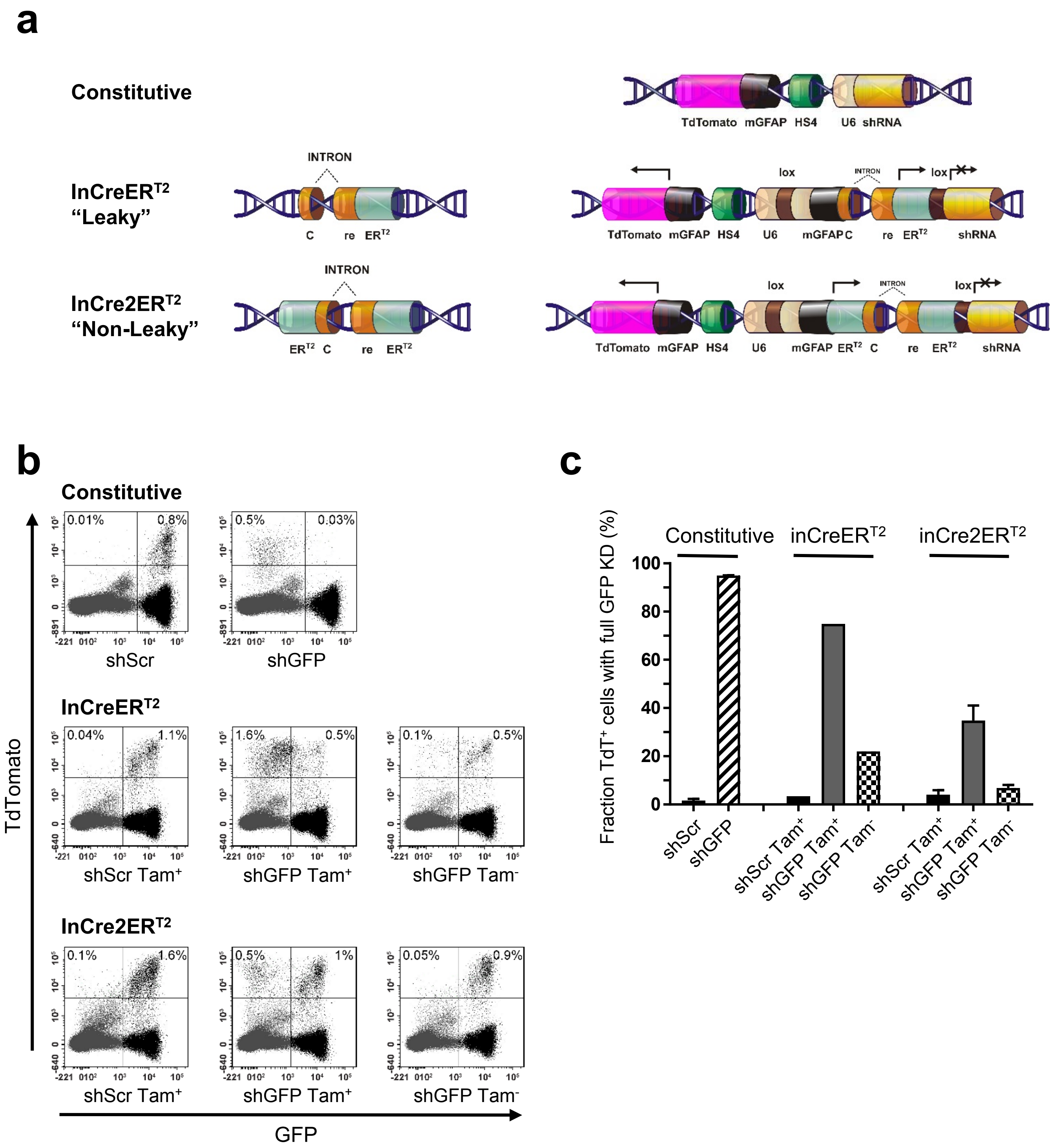


Slezak, de Vin et al., Supplementary Figure 4
