## Supplementary material for "Flexible, fast and selective genetic manipulation of the vertebrate CNS with misPiggy"

### EXPERIMENTAL PROCEDURES

#### Animal experiments.

*Mouse and rat experiments:* All animal procedures were performed in accordance with the regulations of the Institutional Animal Care and Use Committee of KU Leuven. Animals were kept in a specified pathogen free (SPF) facility with controlled humidity, temperature and light conditions. Water and food were provided *ad libitum*.

*Ferret experiments:* Normally pigmented, sable ferrets (*Mustela putorius furo*) were purchased from Marshall Farms (North Rose, NY, USA). All procedures were performed in accordance with protocols approved by the Animal Care Committee of Kanazawa University.

#### Plasmids.

All plasmids used in this study will be deposited at Addgene, along with full plasmid maps, on manuscript publication. Until that point, plasmids can be obtained under a Material Transfer Agreement (MTA) by contacting Dr. Matthew Holt.

Short hairpin RNAs (shRNAs) were used for gene silencing, as these possess several advantages for rapid and flexible screening of gene function *in vivo*[1]. First, the machinery responsible for RNA interference is present in all mammalian cells and does not require the presence of additional accessory factors. Second, shRNAs target mRNAs, which means that selective gene interference is possible with high quality transcriptome information, which is generally more readily available than the genomic information required for gene editing methodologies (e.g location of compatible protospacer adjacent motif (PAM) sites). Further, experience with the technique has led to the publication of potent and specific shRNA sequences for the majority of (mouse) genes, which will produce a homogeneous phenotype across cells, rather than random mono- or biallelic modifications (which may have variable penetrance across cells). Finally, multiple transcript variants can be targeted by sequence (even when transcripts arise from independent promoters).

***In Utero* Electroporation.**

*Mouse experiments: In utero* electroporation experiments were performed essentially as described[2] with the addition of a third electrode for precise targeting of plasmids to visual cortex[3].

Thin walled borosilicate glass (GCT150T-10, Harvard Apparatus) was pulled into a fine needle using a micropipette puller (Sutter Instruments, Model P-97) and then sharpened to a fine point with a micropipette grinder (Narishige, EG-44). Needles were loaded with plasmid DNA (containing 0.01% Fast Green dye (Sigma F7252) as an injection marker) and connected to a custom made tubing system allowing precise mouth-controlled ejection of DNA.

IUE was performed on pregnant CD1 females or Aldh1l1-EGFP transgenic animals (CD1 background) usually at embryonic day (E)15.5 - as this allowed for more robust embryo handling and larger litter sizes. Pregnant animals were anesthetized by intramuscular injection of a ketamine (180 mg/kg) and xylazine (24 mg/kg) mix. Anesthesia was confirmed by loss of the toe-pinch reflex. Animals were then placed on a heating pad set to 37°C to maintain body temperature and eyes were covered with Duratears ophthalmic ointment to prevent excessive drying. The surgical area was then disinfected with 100% ethanol, carefully shaved with a surgical blade and covered with a piece of sterile gauze (with a hole over the surgical area). A small vertical incision (~ 2 cm) was made with sharp scissors to expose the uterus. The uterus (and other exposed organs) were moistened with sterile saline (preheated to 37°C) to prevent drying during the procedure. Embryos were pulled onto the surrounding gauze using ring forceps and fingers.

Approximately 2 µl of plasmid DNA (at a concentration of approx. 2.5 µg/µl) were administered unilaterally to the lateral ventricle. After visually confirming that the ventricle had been successfully targeted, electroporation was performed. In order to target visual cortex, tweezer electrodes (CUY650P5, Sonidel), connected to the negative pole of an electroporator (ECM830 Square Wave Electroporation System, BTX, Harvard Apparatus), were placed on either side of the embryo's head. A third electrode (custom made from a 5 x 3 x 0.5 mm platinum block glued to a cell scraper), which was connected to the positive pole of the electroporator, was centered above the visual cortex of the injected hemisphere. Five 37 V pulses of 50 ms duration, spaced with 950 ms intervals were applied.

After electroporation of embryos (usually 6-10 per litter), the uterus was placed back in the peritoneum. The peritoneal wall was sutured closed (Vicryl V310H, 4-0 taper point, Johnson & Johnson), while the skin incision was closed with metal clips using a Reflex 7 wound clip applicator (Fine Science Tools). The procedure usually took less than 30 minutes per female. After completion, the fur was dried with a hair dryer to prevent heat escape. Finally, the wound was covered with Fucidine ointment (Leo Pharma) to prevent infection. Mice also received a subcutaneous injection of analgesic (0.25 mg carprofen; Zoetis). Mice were left on the heating pad until they had recovered locomotor activity. Mice were then returned to their home cages. Additional carprofen was provided in the drinking water (0.075 mg/ml) for 48 hours after surgery[4]. Pregnant females proceeded through gestation, pups were delivered naturally and weaned. In all experiments, TdTomato (TdT) was used as a positive marker for electroporation of the cortex and $TdT^+$ animals were easily identified by basic widefield imaging of pups at postnatal day (P)3 or P4 (Figure 1a). $TdT^+$ animals were left to develop to the experimental ages indicated in the Main Text.

*Rat experiments:* IUE was performed on pregnant rats at E16.5 or E17.5. The protocol used was identical to that for mouse with minor modifications. Rats were anesthetized using isofluorane (induction 5%, 2.5 l/min $O_2$; maintenance 2.5%, 2.5 l/min $O_2$) and electroporation pulses were delivered at 50 V.

*Ferret experiments: In utero* electroporation in ferrets was performed according to published protocols[5]. Pregnant ferrets (E33.5) were deeply anesthetized using 2% isoflurane with $N_2O$ and $O_2$. Animals were then placed on a heating pad to maintain their body temperature. The uterine horns were exposed and kept wet by adding drops of saline intermittently.

Approximately 2-5 µl of DNA solution (at a concentration of 5 µg/µl) containing 0.01% Fast Green dye (as an injection marker) was injected into the lateral ventricle of an embryo using a pulled glass micropipette. The head of each injected embryo was placed between tweezer-type electrodes (diameter 5 mm: CUY650P5, NEPA Gene, Japan). Square electric pulses (100 V, 50 ms) were delivered 5 times at 1 s intervals using an electroporator (ECM830, Harvard Apparatus, USA). The wall and skin of the abdominal cavity were sutured and embryos were allowed to develop normally.

**Tamoxifen Administration.**

*Mouse experiments:* A dose of 2 mg tamoxifen was administered for five consecutive days. The route of administration varied, depending on the experiment. In some instances, tamoxifen was administered to the lactating mother starting at P5 or P10 via intraperitoneal (IP) injection. In other cases, administration was via direct IP injection of animals at P21. When timing of induction was not important, tamoxifen was generally administered to the lactating mother, for experimental ease. Tamoxifen (Sigma T5648) was prepared using two different protocols, without any observable differences in the final results. In one protocol, tamoxifen was dissolved in corn oil (Sigma C8267). In the second protocol, tamoxifen was dissolved in 100% ethanol and this solution was diluted with corn oil (Sigma C8267) at a 1:9 ratio. The final stock concentration of tamoxifen was 10 mg/ml or 20 mg/ml, respectively. The drug was administered by a single intraperitoneal injection of 100 or 200 µl solution (giving a final total dose of 2 mg tamoxifen per day). Tamoxifen was freshly made for each independent set of experiments and was kept at 4°C until injections ceased[6]. Tissues were typically harvested 14 days after the last tamoxifen injection.

*Rat experiments:* A dose of 7 mg tamoxifen was administered for 5 consecutive days to lactating females starting at P5, using intraperitoneal injections[7]. Tamoxifen (Sigma T5648) was dissolved in corn oil (Sigma C8267) at a concentration of 20 mg/ml and administered by injection of 350 µl solution.

*Ferret experiments:* Tamoxifen (Sigma T5648) was dissolved in 100% ethanol and this solution was diluted with corn oil (Sigma C8267) at a 1:9 ratio. 2 mg tamoxifen were administered to ferret pups (P5) by intraperitoneal injection, alternating once or twice per day for 5 consecutive days, starting with a single injection on the first day. Fresh tamoxifen was used for each set of injections. The tamoxifen stock was kept shielded from light at 4°C. Tissues were harvested 14 days after the last tamoxifen injection.

**Histology.**

*Mouse and rat experiments*

- *Tissue fixation and slice preparation:* Animals were administered an overdose of Nembutal (60 mg/kg), before being transcardially perfused with 4% paraformaldehyde (PFA). Brains were then removed and post-fixed overnight in 4% PFA. 50 µm thick sections were then cut using a Leica VT1000S vibratome. Where imaging of TdT alone was sufficient, slices were mounted directly onto slides using Vectashield mounting media (Vectorlabs).

- *Immunostaining:* When antibody-based detection methods were used, sections were routinely permeabilized and blocked for 1 hour at room temperature in 0.2% Triton X-100/PBS supplemented with 10% normal goat serum (NGS). Sections were incubated with primary antibodies (diluted in 0.2% Triton X-100/PBS with 1% NGS) overnight at 4°C. Sections were then given three 10 minutes washes in PBS, before incubation with secondary antibodies (diluted in 0.2% Triton X-100/PBS) for 2-3 hours at room temperature. Sections were then given three 10 minutes washes in PBS, before mounting using Vectashield (Vectorlabs). Primary antibodies used include rabbit anti-GFAP (1:300 dilution, Dako, Z0334), rabbit anti-S100β (1:300 dilution, Synaptic Systems, 287003), mouse anti-APC (1:200 dilution, Abcam, ab16794), rat anti-PDGFRα (1:600 dilution, BD Pharmingen, 558774), rabbit anti-Iba1 (1:600 dilution, Synaptic Systems, 234003), guinea pig anti-NeuN (1:250 dilution, Synaptic Systems, 266004-1) and rabbit anti-GFP (1:300 dilution, Synaptic Systems, 132002). Secondary antibodies used include goat anti-rat Alexa Fluor 488 (Abcam, ab150157), goat anti-guinea pig Alexa Fluor 488 (Jackson Immuno, 106-545-003), goat anti-rabbit Alexa Fluor 488 (Jackson

Immuno, 111-545-144), and goat anti-mouse Alexa Fluor 488 (Jackson Immuno, 115-545-146). All secondary antibodies were used at a dilution of 1:400.

*Ferret experiments*

- *Tissue fixation and slice preparation:* Animals were deeply anesthetized with 10% pentobarbital and transcardially perfused with 4% PFA. Dissected brains were further post-fixed overnight with 4% PFA. To make coronal sections, the brains were cryoprotected by immersion for three days in 30% sucrose (in PBS) and embedded in OCT compound (Sakura). Sections of 50 µm thickness were prepared using a cryostat (Leica CM1850).

- *Immunostaining:* Sections were permeabilized with 0.3% Triton X-100 in PBS and blocked with 2% skimmed milk, 0.3% Triton X-100 in PBS. The sections were incubated overnight with rabbit anti-GFP antibody (1:300 dilution, Medical & Biological Laboratories, Japan, 598). After being washed, sections were incubated with Alexa Fluor 488-conjugated donkey anti-rabbit IgG antibody (1:500 dilution, Thermo Fisher Scientific, A21206) and 1 µg/ml Hoechst 33342 (Thermo Fisher Scientific). Sections were then washed and mounted.

**Image analysis.**

*Mouse Experiments*

**-** *Quantification of IUE efficiency using nls-TdT expression:* P21 brains were perfused and 50 µm vibratome sections were cut. Coronal tissue sections covering the whole brain were then mounted from front to back (at approximate 550 µm intervals) and imaged using a Zeiss Slide scanner (Axio Scan Z1) with a PL APO 20x/0.8 objective to pinpoint the sections expressing TdT. As expected, TdT expression was highly focused in the brain and present over a limited number of consecutive brain sections, consistent with focal electroporation and plasmid uptake. The number of $TdT^+$ cells in these sections was used as a relative measure of electroporation efficiency. For each brain analyzed, three sections with the highest expression density were selected for further analysis. In this case, expression density was calculated as the number of $TdT^+$ cells per unit area. TdT expression was usually well demarcated, meaning that the measured areas could simply be set using the boundaries of TdT expression (Figure 1b, Supplementary Figure 2). Confocal images were acquired on a Leica SP8 microscope equipped with a HC PL APO CS2 20x/0.70 objective using a 0.75x zoom setting. Image acquisition was controlled using Las X v2.0 software. Z-stacks were taken in the tissue sections with the highest expression density. Each Z-stack was 10 μm total thickness. The thickness of each image in the stack was 0.79 μm, with x-y dimensions of 775 μm x 775 μm. For each tissue section, more than 50% of the transfected area was imaged (with great care taken not to scan different fields of view that overlapped). Following acquisition, Leica imaging files were imported into Image J. Maximum projection images were made for both the GFP and TdT channels. To determine the efficiency of transfection, the maximum projection images for each color channel were merged and the number of GFP labelled cells (astrocytes) containing TdT labelled nuclei was determined, as a measure of the efficiency of transfection (Supplementary Figure 2). Data were generally collected from three different brains (typically three sections per brain), with over 9000 $GFP^{+}$ cells checked for colocalization with nls-TdT signal for each experimental condition analyzed.

- *Fluorescence intensity measurements:* Brains were sliced and sections were selected based on high levels of TdT fluorescence, as described for the quantification of IUE efficiency. Once sections were identified, images were acquired using a Zeiss Slide scanner with a PL APO 20x/0.8 objective (see above). Acquisition conditions were kept constant across sections. To quantify fluorescence intensity using ZEN analysis software (Zeiss), three regions of interest (ROIs) measuring 630 μm x 630 μm were placed in the ipsilateral hemisphere, evenly distributed across the area of TdT fluorescence (Figure 1b, Supplementary Figure 3). As a control, three boxes of equivalent size were placed in the contralateral (non-electroporated) hemisphere, directly mirroring their position in the ipsilateral hemisphere. The fluorescence intensity (in arbitrary units) was measured in each box for GFAP and IBA1 staining. Data are expressed as the ratio between the ipsilateral and contralateral measurements.

- *Analysis of IBA1⁺ cell density:* Confocal Z-stack images (0.79 µm section thickness, 10 µm total depth, x-y dimensions 775 µm x 775 µm) were taken in the ROIs used for the analysis of GFAP and IBA1 fluorescence intensities (see above). Images were acquired on a Leica SP8 microscope equipped with a HC PL APO CS2 20x/0.70 objective at 0.75x zoom. Maximum projection images of the Z-stacks were made and cells were manually counted using the ImageJ cell counter plug in. Data are expressed as the average number of cells per unit area in the ipsilateral or contralateral hemispheres.

*Ferret experiments:* Confocal images were acquired on a Leica SP8 microscope equipped with a HC PL APO CS2 20x/0.70 objective using a 1.5x zoom setting. Image acquisition was controlled using Las X v2.0 software.

***In vivo* 2-photon imaging.**

*Cranial window insertion:* Cranial windows were inserted following published protocols[8]. Mice were administered intramuscular dexamethasone (4 mg/kg) at least 4 hours before surgery to prevent neural edema both during and after the craniotomy. Anesthesia during surgery was using isofluorane (3% isoflurane in 100% oxygen at a flow rate of 0.8 l/min for induction, 1.5% isofluorane in 100% oxygen at a flow rate of 0.5 l/min for maintenance).

Following induction of anesthesia (checked using the toe pinch reflex), eyes were covered with Duratears ophthalmic ointment (Alcon). The fur on the skull was shortened with scissors and the remainder removed using a depilating gel. The area was then cleaned using 70% ethanol and iodine solution to minimize the risk of infection. The scalp was removed using Dumont forceps and fine scissors, the periosteum was cleared with a scalpel blade and the temporalis muscle separated from the skull. A custom-made titanium head plate[9] was centered to the posterior left hemisphere and attached to the skull with cyanoacrylic glue. The exposed muscles and skull were covered with a thin layer of Vetbond (3M) and dental cement (C&B Superbond, Générique International). A 5 mm diameter craniotomy was performed centered on the visual cortex (1.6 mm anterior to lambda, 3.1 mm lateral to midline). Bone was removed with forceps and the underlying dura was left intact. The craniotomy was covered with a window consisting of an 8 mm diameter cover glass (orientated to the top) glued with optical adhesive (NOA71, Norland) to two 5 mm diameter cover glasses. This window was then fixed to the skull using acrylic glue and dental cement (Kett Tab 2000, KemDent). A custom well (to hold water for the immersion objective) was formed using neoprene O-rings. After surgery, mice were left on a 37°C heating pad until they had recovered locomotor activity. Mice were treated with cefazoline (100 mg/kg) and buprenorphine (0.04 mg/kg) immediately after recovery and twice daily for three consecutive days.

*Training:* Mice were habituated to head fixation before imaging sessions[10]. Training was started a minimum of six days following cranial window implantation. During training, cage housed mice had restricted access to water (1 ml per day), unless mice showed significant weight loss (>15% of initial weight pre-surgery), in which case water was available *ad libitum*. During the first five days, mice were habituated to handling and head fixation. In the following days (in sessions up to 30 min duration), mice were head-fixed on a treadmill platform and were trained to run on a treadmill belt 130 cm in length. Mice were rewarded with tap water or 10% sucrose solution after completing a lap on the treadmill (5-10 µl drop size).

*Imaging:* During each imaging session, mice were head-fixed and allowed to move on the treadmill belt as during the training sessions, with a water or sucrose reward given after each completed lap. Images of GCaMP6m fluorescence were obtained from layer 2/3 astrocytes in the visual cortex (125 to 300 µm depth) using a custom-designed microscope (Neurolabware), equipped with a Nikon objective (16x/0.8). The microscope was controlled by a computer running custom-written software (Neurolabware). Two photon excitation was achieved using a MaiTai DeepSee laser corrected for group-delay dispersion. The laser wavelength was tuned to 920 nm and scanned across the field of view at a rate of 31 frames per second (fps) using a galvo-based system. Typically, an area roughly equal to 620 mm x 380 mm of the visual cortex was imaged (512 lines of 1154 pixels). Laser power was limited to 20 to 60 mW at the objective, depending on the depth of the imaging plane, to minimize phototoxicity. Emitted light was collected using appropriate filters (green fluorescence: 510 nm/584 nm; red fluorescence: 630 nm/692 nm; Semrock) and GaAsP photomultiplier tubes (Hamamatsu).

*Analysis of data from in vivo imaging experiments:* Raw images were reconstructed and corrected for motion artefacts using custom MATLAB routines. Regions of interest (ROIs) were manually superimposed on the images, outlining astrocytes based on morphology. GCaMP6m response to $Ca^{2+}$ binding was calculated as the average increase in GCaMP intensity relative to baseline fluorescence (dF/F) for each individual ROI.

**Flow cytometry experiments.**

*Tissue dissociation:* Mice were sacrificed by cervical dislocation at P21 in the case of experiments looking at constitutive expression of shRNA, or at P30 in the case of inducible expression of shRNA. CNS tissue was recovered and the electroporated region localized using an Olympus SZX16 fluorescence microscope with appropriate filters for TdT (Figure 1a). This region was then dissected out in ice-cold Hank's Balanced Salt Solution (HBSS) without calcium and magnesium (Sigma). A single cell suspension was then prepared using the Tissue Dissociation Kit P (Miltenyi Biotec), with minor modifications[11]. Briefly, following 35 minutes of enzymatic digestion, the tissue was mechanically dissociated (3 rounds of trituration, 10 strokes each round) with 5 ml serological pipettes. Freshly dissociated cells were then passed through a 20 µm Nitex filter (SEFAR) to remove remaining tissue clumps. Myelin removal using equilibrium density centrifugation was then performed: 90% Percoll PLUS (Life Sciences) in 1x HBSS with calcium and magnesium (Sigma) was added to the cell suspension to give a final Percoll concentration of 24%. DNaseI (Worthington) was then added (1250U per 10 ml of suspension), the suspension was mixed and then spun down at $300g_{Av}$ for 11 min at room temperature in a Hettich 320R universal centrifuge (with minimal brake). The cell-containing pellet was resuspended in 0.5% BSA (Sigma) in phosphate buffered saline (PBS) without calcium and magnesium (Thermo Fisher Scientific) and filtered through a 20 µm Nitex mesh.

*Flow cytometry:* Flow cytometry was performed as previously described[11], using a BD FACSCanto I (BD Biosciences), controlled using FACSDiva software (version 6.1.3). Cells were initially gated using the forward-scatter (FSC) versus side scatter (SSC) plot, with doublets being discriminated using the FSC-width/FSC-area plot. Compensations were done using single-stained controls and setting the median fluorescence of positive and negative populations at the same level. Gates were set on unstained controls or fluorescence minus one (FMO) controls. The viability dye 7-AAD (eBioscience) was used to discriminate dead cells.

**Supplemental Figure and Table Legends.**

**Supplementary Figure 1. Cell type specific expression is critical for precise genetic modifications of cell function**

(**a**) Astrocyte specific expression using a modified GFAP promoter (mGFAP). Representative images and orthogonal projections from brain slices of an adult mouse (P21) electroporated at E16.5 with a construct constitutively expressing TdTomato (TdT). Cells possessed astrocyte-like morphology. Slices were counter-stained with antibodies against cell-type markers: GFAP and S100β (astrocytes), NeuN (neurons), APC (oligodendrocytes), IBA1 (microglia), and PDGFRα (NG2+ cells). Scale bars, 20 μm.

(**b**) Neuron specific expression using a human synapsin 1 (hSYN1) promoter. Representative images of brain slices obtained from an adult mouse (P40) electroporated at E15.5. Constitutive expression of TdT in astrocytes was driven by a mGFAP promoter. The hSYN1 promoter was used to drive expression of tamoxifen inducible Cre. Recombination led to expression of the genetically encoded calcium indicator GCaMP6m in cells with typical neuronal morphology. Images are from animals treated with tamoxifen ($Tam^+$) or with vehicle ($Tam^-$). Scale bars, 100 μm.

(**c**) Oligodendrocyte specific expression using a proteolipid protein (PLP) promoter. Representative images of brain slices obtained from an adult mouse (P50) electroporated at E15.5. Constitutive expression of TdT in astrocytes was driven by a mGFAP promoter. The PLP promoter was used to drive expression of tamoxifen inducible Cre. Following tamoxifen, recombination led to expression of the genetically encoded calcium indicator GCaMP6m in cells with typical oligodendrocyte morphology, which were found in the corpus callosum. Images are from animals treated with tamoxifen (Tam$^{+}$) or with vehicle (Tam$^{-}$). Scale bars, 100 μm.

Data are representative images seen in 3 slices from 3 independent mice electroporated with each construct.

**Supplementary Figure 2. Efficiency of the electroporation system.**

(**a**) Two constructs were used to test the efficiency of the electroporation system. Both constructs used TdTomato (TdT) fused to a nuclear-localization signal (nls), as this allowed unambiguous cell identification. The first construct expressed nls-TdT under control of a modified (shortened) GFAP promoter (mGFAP), which was used to investigate the effect of developmental age (E14.5 versus E16.5) on electroporation efficiency. The total size of the plasmid was 10.6 kb, with a transgene cassette of 2.6 kb. The second construct was identical to the first except for the insertion of the full-length mouse GFAP promoter into the transgene cassette (increasing plasmid size to 21.3 kb with a transgene cassette of 13.3 kb). This was done to evaluate how increasing the size of the transgene cassette, and consequently plasmid size, influences electroporation efficiency at E16.5. Representative images of brain slices obtained from an adult Aldh1l-EGFP mouse (P21) electroporated at E16.5 with the 10.6 kb plasmid. Scale bar, 30 μm.

(**b**) Summary of data for the conditions tested. Efficiency is the fraction of GFP+ cells expressing TdT. More than 9,000 GFP+ cells were assessed for nls-TdT expression in each condition, over a minimum of three experimental animals per condition. Data are the average ± SD. * $P<0.05$, ** $P<0.01$ (Unpaired *t*-test)

**Supplementary Figure 3. Lack of gliosis in the adult brain after *in utero* electroporation.**

(a) Left: Low magnification images of brain slices from an adult mouse (P21) electroporated with a construct constitutively expressing TdT in astrocytes at E16.5 (ipsilateral side). Slices were stained using antibodies against GFAP (reactive astrocytes) and IBA1 (microglia), as markers for reactive gliosis induced by the electroporation procedure. Right: Average fluorescence across the indicated regions (boxes). Values were normalized to those measured on the non-electroporated (contralateral) side. Scale bars, 500 µm. Data are from 9 slices from 3 independent mice and are given as average ± SD.

(b) Left: High magnification images from the visual cortex of electroporated brains stained for microglia using IBA1. Right: Average number of microglia measured per unit area in both ipsilateral and contralateral sections. Scale bars, 30 µm. Data are from 9 slices from 3 independent mice and are given as average ± SD.

**Supplementary Figure 4. Development of tamoxifen inducible systems.**

(a) Ligand inducible constructs were created by fusing one or two tamoxifen response elements ($ER^{T2}$) to intronized Cre (inCre), creating $inCreER^{T2}$ and $inCre2ER^{T2}$, respectively. Right: Strict ligand inducibility was tested using shRNA mediated knock down of GFP as a readout. The constructs used in experiments contained either a constitutively active U6 promoter or a Cre-activated (split) U6 promoter. Cre expression was limited to astrocytes using a specific mGFAP promoter. Constitutive expression of TdT was used as a control for successful electroporation.

(b) Constructs allowing constitutive or inducible expression of shRNA against GFP (or a scrambled control) were electroporated into Aldh1l1-EFGP mice at E15.5. When appropriate, animals received tamoxifen ($Tam^+$) or vehicle control ($Tam^-$) at P5-P9 via the lactating mother. Single cell suspensions of brain were produced, which were then subjected to flow cytometry analysis. Figures represent the proportion of cells in each particular gate.

(c) Summary of flow cytometry data obtained with the various constructs. Columns show the percentage of $TdT^+$ cells showing *complete* loss of GFP (which likely underestimates knock down efficiency across the cell population). The efficiency of gene knock down using a constitutively expressed shRNA against GFP matched or exceeded that found with IUE-based CRISPR-Cas9 systems[12, 13]. The proportion of cells showing knock down using inducible systems was largely influenced by recombination efficiency. Strict ligand inducibility ('non-leaky') was only achieved with the addition of two $ER^{T2}$ domains to Cre (compare to the low level of non-ligand induced ('leaky') knock down seen with one $ER^{T2}$ domain). Conversely, recombination efficiency was lowest with the $inCre2ER^{T2}$ system, as expected due to more efficient sequestration of inCre by cytosolic chaperones[14].

**Supplementary Table 1. Detailed description of constructs used in the study.**

Individual expression modules are listed. MP, misPiggy; CS, cloning site. Constructs are listed sequentially.

**Supplementary References.**

1. Boettcher, M. & McManus, M.T. Choosing the Right Tool for the Job: RNAi, TALEN, or CRISPR. *Mol Cell* **58**, 575-585 (2015).

2. Tabata, H. & Nakajima, K. Efficient *in utero* gene transfer system to the developing mouse brain using electroporation: visualization of neuronal migration in the developing cortex. *Neuroscience* **103**, 865-872 (2001).

3. dal Maschio, M. et al. High-performance and site-directed *in utero* electroporation by a triple-electrode probe. *Nat Commun* **3**, 960 (2012).

4. Ingrao, J.C. et al. Aqueous stability and oral pharmacokinetics of meloxicam and carprofen in male C57BL/6 mice. *J Am Assoc Lab Anim Sci* **52**, 553-559 (2013).

5. Kawasaki, H., Iwai, L. & Tanno, K. Rapid and efficient genetic manipulation of gyrencephalic carnivores using *in utero* electroporation. *Mol Brain* **5**, 24 (2012).

6. Madisen, L. et al. A robust and high-throughput Cre reporting and characterization system for the whole mouse brain. *Nat Neurosci* **13**, 133-140 (2010).

7. Schönig, K. et al. Conditional gene expression systems in the transgenic rat brain. *BMC Biol* **10**, 77 (2012).

8. Goldey, G.J. et al. Removable cranial windows for long-term imaging in awake mice. *Nat Protoc* **9**, 2515-2538 (2014).

9. Andermann, M.L., Kerlin, A.M. & Reid, R.C. Chronic cellular imaging of mouse visual cortex during operant behavior and passive viewing. *Front Cell Neurosci* **4**, 3 (2010).

10. Andermann, M.L. et al. Chronic cellular imaging of entire cortical columns in awake mice using microprisms. *Neuron* **80**, 900-913 (2013).

11. Batiuk, M.Y. et al. An immunoaffinity-based method for isolating ultrapure adult astrocytes based on ATP1B2 targeting by the ACSA-2 antibody. *J Biol Chem* **292**, 8874-8891 (2017).

12. Chen, F., Rosiene, J., Che, A., Becker, A. & LoTurco, J. Tracking and transforming neocortical progenitors by CRISPR/Cas9 gene targeting and piggyBac transposase lineage labeling. *Development* **142**, 3601-3611 (2015).

13. Shinmyo, Y. et al. CRISPR/Cas9-mediated gene knockout in the mouse brain using *in utero* electroporation. *Sci Rep* **6**, 20611 (2016).

14. Matsuda, T. & Cepko, C.L. Controlled expression of transgenes introduced by *in vivo* electroporation. *Proc Natl Acad Sci U S A* **104**, 1027-1032 (2007).
